# Genetic and phenotypic differentiation of lumpfish (*Cyclopterus lumpus*) across the North Atlantic: implications for conservation and aquaculture

**DOI:** 10.1101/382119

**Authors:** Benjamin Alexander Whittaker, Sofia Consuegra, Carlos Garcia de Leaniz

**Affiliations:** Centre for Sustainable Aquatic Research Swansea University Singleton Park, Swansea, SA2 8PP, Wales, UK

**Keywords:** translocation, conservation genetics, migration, aquaculture, cleaner fish

## Abstract

Demand for lumpfish (*Cyclopterus lumpus*) has increased exponentially over the last decade, both for their roe, which is used as a caviar substitute, and increasingly also as cleaner fish to control sea lice in salmon farming. The species is classified as Near Threatened by the UICN and there are growing concerns that over-exploitation of wild stocks and translocation of hatchery-reared lumpfish may compromise the genetic diversity of native populations. We carried a comparative analysis of genetic and phenotypic variation across the species’ range to estimate the level of genetic and phenotypic differentiation, and determined patterns of gene flow at spatial scales relevant to management. We found five genetically distinct groups located in the West Atlantic (USA, and Canada), Mid Atlantic (Iceland), East Atlantic (Faroe Islands, Ireland, Scotland, Norway, and Denmark), English Channel (England) and Baltic Sea (Sweden). Significant phenotypic differences were also found, with Baltic lumpfish growing more slowly, attaining a higher condition factor and maturing at a smaller size than North Atlantic lumpfish. Estimates of effective population size were consistently low across the NE Atlantic (Iceland, Faroe Islands, Norway), the area where most wild lumpfish are fished for their roe, and also for the aquaculture industry. Our study suggests that some lumpfish populations are very small and have low genetic diversity, which makes them particularly vulnerable to over-exploitation and genetic introgression. To protect them we advocate curtailing fishing effort, closing the breeding cycle of the species in captivity to reduce dependence on wild stocks, restricting the translocation of genetically distinct populations, and limiting the risk of farm escapes.

## 1 INTRODUCTION

The control of parasitic sea-lice (*Lepeophtheirus salmonis*) is one of the most pressing problems facing salmon farming (Torrissen et al. 2013; Treasurer 2002), as sea-lice have become resistant to chemical treatment (Aaen et al., 2015; Lees et al., 2008) and threaten the sustainability of the industry. Several species of cleaner fish have been used as an alternative to the use of antiparasitic therapeutants (Treasurer 2018), but the lumpfish (*Cyclopterus lumpus*) is probably the most useful as, in contrast to other cleaner fish like wrasse, it continues to feed on sea lice at low temperatures and is easy to rear in captivity (Imsland et al., 2014; Powell et al., 2017). Demand for lumpfish has increased exponentially since 2012 (Powell et al., 2017; Treasurer, 2018). However, nearly all lumpfish used in salmon farming are still derived from wild broodstock (Jonassen et al., 2018a), and as they are generally used in a single salmon production cycle (Powell et al., 2017), satisfying aquaculture demands can put considerable pressure on wild stocks.

Lumpfish has been classified as Near Threatened in the IUCN Red List (Lorance et al. 2015), but information on the conservation status of different populations is very limited, and it is likely that some populations are already overexploited (Myers & Sjare, 1995; Powell et al., 2017). Ripe females have traditionally been targeted for their roe, which is processed and sold as a cheap alternative to caviar, and while the Icelandic and Greenland lumpfish fisheries are closely monitored, others are largely unregulated (Powell et al., 2017; Kousoulaki et al., 2018). A strong reduction in catch per unit effort has been detected in some lumpfish fisheries over the last 25 years (Lorance et al., 2015), and there are concerns that removing additional spawners for the expanding lumpfish aquaculture industry could impact on some small populations (Hedeholm et al., 2014; Powell et al. 2017, Powell et al., 2018), as it has been reported for other cleaner fish fisheries (Halvorsen et al., 2017).

Stock movements represent an additional risk to wild lumpfish as large numbers of hatchery-reared lumpfish are being translocated across the North Atlantic to supply salmon farms (Jonassen et al., 2018b; Treasurer et al., 2018) and this could pose a potential threat to local populations. For example, over 85% of all lumpfish deployed in Scotland during 2017 originated from eggs imported from Iceland and Norway, and none came from local sources (Treasurer et al., 2018). In Ireland, 70% of lumpfish deployed during 2015-2016 were derived from eggs imported from Iceland and Norway (Bolton-Warberg et al., 2018), while in the Faroe Islands nearly all lumpfish used during 2014-2016 were of Icelandic origin (Steinarsson and Arnason, 2018; Johanssen et al., 2018). There is a danger that if non-native lumpfish escape from salmon farms they could interbreed with local populations and result in genetic introgression (Powell et al., 2017), as has been reported for farmed salmonids (e.g. Consuegra et al., 2011). Lumpfish translocations are likely to intensify in the near future (Jónsdóttir et al., 2017), and while escapes of lumpfish have not yet been reported, these seem largely inevitable in open salmon net-pens, as have already been documented for two species of wrasse (Jansson et al., 2017; Faust et al., 2018). Whether escapes can have a genetic impact on local lumpfish populations will depend on the number of escapees, their reproductive success, and the extent of genetic differentiation between local and introduced fish, but none of these parameters are currently known.

Lumpfish are distributed across a vast marine area, extending to both sides of the North Atlantic and into the Baltic (Davenport 1985; Powell et al., 2018), and there is thus scope for substantial differentiation. Soon after hatching, the larvae attach to the substrate using a specialized suction cup, which probably limits larval dispersal (Davenport, 1985). Tagging studies suggest that, although adults can swim up to 49 km/day, some individuals remain within a restricted 80 km range after +250 days at liberty (Kennedy et al., 2015). There is also evidence of homing (Kennedy et al., 2014), which will favour reproductive isolation and may result in stock differentiation. For example, spawning time may vary by two months within single populations (Wittwer and Treasurer, 2018), but as much as seven months among populations, from January in the English channel (Powell et al., 2018) to August near the Arctic circle (Jónsdóttir et al., 2018). Population differences also exist in growth and delousing behaviour (Johannesen et al., 2018) and, as these are maintained under common rearing conditions (Imsland et al., 2016; Bolton-Warberg et al., 2018), they are likely to be inherited. Such differences suggest that lumpfish may form discrete populations, and that these may be adapted to local conditions. Yet, the extent of genetic differentiation in lumpfish is uncertain. Thus, while significant genetic differences have been found at large spatial scales (i.e. Canada vs Norway; Pampoulie et al., 2014), populations at smaller scales appear to be relatively homogenous. For example, lumpfish sampled in the English channel appear to be to largely undifferentiated (Consuegra et al., 2015), as do fish sampled along the Norwegian coast (Jónsdóttir et al., 2017). In contrast, in Greenland two genetically distinct groups have been found in the north and south (Garcia-Mayoral et al., 2016) suggesting that there can also be some fine scale genetic structuring.

### 1.1 Aims

The aims of our study were three. First, given their limited larval dispersal and evidence for homing, we hypothesised that lumpfish might display genetic isolation by distance, with populations closer together being more genetically similar than those further apart (Rousset 1997). By sampling across the whole range, we aimed to estimate the level of genetic and phenotypic differentiation, and determine the patterns of gene flow across the species’ range, at spatial scales relevant to management.

Secondly, given the level of recent stock transfers, we wanted to know to what extent translocations could pose a potential genetic risk to local populations. For this, we provide genetic baselines on wild populations against which genetic introgression from farm escapees can be latter assessed, as done for Atlantic salmon (Gilbey et al., 2017).

Finally, as some lumpfish populations may be endangered, we provide estimates of effective population size, and test for the existence of genetic bottlenecks to better understand their conservation status and the extent to which gene flow could mitigate the impacts of over-exploitation.

## 2 MATERIAL AND METHODS

### 2.1 Collection of samples

Fin tissue was obtained from 410 lumpfish originating from 15 sites across the species’ range (Table 1) and were stored in 96% ethanol at −20°C until analysis. Sites located within an 80 km radius (the estimated maximum range of dispersal, Kennedy et al., 2015) were pooled together to minimise the risk of spatial pseudo-replication. Samples were pooled from the Faroe Islands (Klasvík and Kollafjørður, c. 20km), Denmark (Køge Bay and Mosede Havn, c. 13km) and Sweden (survey hauls from Bornhölm to Öland, and from Gotland to Gotska Sandön). Pooled groups were named after the site contributing the largest number of samples. Biometric data on length (mm) and weight (g) were available for eight of the 15 sites (Table 1).

**Table 1.** Details of study sites sampled for lumpfish (*N*: sample size for genetic analysis; *Nb*: sample size for biometric analysis; *N_A_* = mean number of alleles (± SE), *N_E_* = mean number of effective alleles, *N_PA_* = number of private alleles, *H_O_* = observed heterozygosity, *H_E_* = expected heterozygosity, *F_IS_* = fixation index; *denotes deviation from HWE due to heterozygote deficiency after Bonferroni correction at *P*<0.0033

| Year | Country | Site | Lat. | Long. | $N$ | $Nb$ | | $N_A$ | $N_E$ | $N_{PA}$ | $H_O$ | $H_E$ | $F_{IS}$ |
| --- | --- | --- | --- | --- | --- | --- | --- | --- | --- | --- | --- | --- | --- |
| 2016 | USA | Frenchman Bay (FB) | 44.33 | -68.15 | 30 | - | mean<br>± SE | 6.0<br>0.775 | 3.100<br>0.548 | 0.300<br>0.213 | 0.566<br>0.058 | 0.613<br>0.044 | 0.100<br>0.050 |
| 2016 | USA | Cobscook Bay (CB) | 44.90 | -67.05 | 30 | - | mean<br>± SE | 6.1<br>0.862 | 3.452<br>0.456 | 0.100<br>0.100 | 0.640<br>0.063 | 0.668<br>0.038 | 0.078<br>0.055 |
| 2016 | Canada | Witless Bay (WB)* | 47.21 | -52.69 | 30 | 30 | mean<br>± SE | 6.7<br>0.870 | 3.459<br>0.425 | 0.400<br>0.163 | 0.630<br>0.049 | 0.673<br>0.036 | 0.080<br>0.050 |
| 2016 | Iceland | Hafnir (Ha) | 63.93 | -22.69 | 30 | - | mean<br>± SE | 5.5<br>0.500 | 2.971<br>0.222 | 0.000<br>0.000 | 0.637<br>0.041 | 0.643<br>0.031 | 0.019<br>0.050 |
| 2016 | Faroe Is. | Klasvík (KI)* | 62.23 | -6.58 | 30 | - | mean<br>± SE | 6.8<br>0.359 | 3.713<br>0.453 | 0.200<br>0.133 | 0.668<br>0.049 | 0.700<br>0.030 | 0.065<br>0.047 |
| 2014 | Ireland | Ventry Bay (VB) | 52.20 | -10.12 | 30 | 26 | mean<br>± SE | 6.8<br>0.389 | 3.255<br>0.346 | 0.100<br>0.100 | 0.647<br>0.056 | 0.658<br>0.038 | 0.032<br>0.050 |
| 2017 | Scotland | Outer Hebrides (OH) | 58.16 | -6.38 | 30 | 18 | mean<br>± SE | 6.5<br>0.453 | 3.247<br>0.448 | 0.000<br>0.000 | 0.623<br>0.055 | 0.644<br>0.041 | 0.060<br>0.036 |
| 2015 | England | Weymouth (We) | 50.61 | -2.46 | 30 | 30 | mean<br>± SE | 5.8<br>0.593 | 2.979<br>0.423 | 0.000<br>0.000 | 0.607<br>0.062 | 0.597<br>0.059 | -0.012<br>0.053 |
| 2015 | England | Guernsey (Gu) | 49.47 | -2.59 | 30 | 30 | mean<br>± SE | 5.6<br>0.476 | 3.068<br>0.431 | 0.000<br>0.000 | 0.618<br>0.083 | 0.608<br>0.055 | 0.032<br>0.084 |
| 2017 | Norway | Namsen (Na) | 59.15 | 6.01 | 21 | 21 | mean<br>± SE | 6.3<br>0.539 | 3.080<br>0.470 | 0.100<br>0.100 | 0.576<br>0.076 | 0.614<br>0.050 | 0.105<br>0.078 |
| 2016 | Norway | Averøy (Av) | 63.05 | 7.48 | 30 | - | mean<br>± SE | 5.7<br>0.496 | 3.077<br>0.336 | 0.000<br>0.000 | 0.677<br>0.051 | 0.638<br>0.038 | -0.038<br>0.030 |
| 2015 | Norway | Rogaland (Ro) | 64.45 | 11.41 | 19 | - | mean<br>± SE | 4.5<br>0.342 | 2.625<br>0.291 | 0.000<br>0.000 | 0.600<br>0.071 | 0.594<br>0.026 | -0.056<br>0.046 |
| 2012 | Denmark | Køge Bay (KB)* | 55.46 | 12.18 | 30 | - | mean<br>± SE | 5.7<br>0.423 | 3.168<br>0.273 | 0.000<br>0.000 | 0.626<br>0.036 | 0.660<br>0.033 | 0.067<br>0.029 |
| 2017 | Sweden | Öland (Öl)* | 55.72 | 16.39 | 16 | 16 | mean<br>± SE | 4.7<br>0.423 | 2.838<br>0.320 | 0.100<br>0.100 | 0.548<br>0.077 | 0.592<br>0.061 | 0.110<br>0.073 |
| 2017 | Sweden | Gotska Sandön (GS)* | 57.95 | 18.97 | 24 | 24 | mean<br>± SE | 5.3<br>0.448 | 3.192<br>0.449 | 0.300<br>0.213 | 0.481<br>0.073 | 0.611<br>0.064 | 0.241<br>0.081 |

### 2.2 DNA extraction and amplification

DNA was extracted using the Nexttec Isolation kit (Nexttec, UK) following the manufacturer’s protocol. The concentration of extracted DNA was quantified using a Nanodrop 2000 (Thermo Fisher Scientific Inc., USA) and diluted with DNA free water to 50ng/μl where necessary. A 2µl of sample DNA was used for amplification using a QIAGEN Multiplex PCR kit (QIAGEN, UK) in a total reaction volume of 9µl. Ten microsatellite loci (*Clu*29, *Clu*34, *Clu*36, *Clu*45 and *Clu*12, *Clu*26, *Clu*33, *Clu*37, *Clu*40, *Clu*44 (Skirnisdottir et al., 2013) were genotyped in two separate multiplex reactions (Table S1). Amplification consisted of a single initial activation step at 95°C for 15 minutes followed by eight cycles of touchdown PCR denaturation at 94°C for 30 seconds, annealing from 64°C or 60°C to 56°C in descending two-cycle steps of 2°C and an extension at 72°C for 90 seconds, 24 additional cycles with an annealing temperature of 56°C and a single final extension at 60°C for 30 minutes. An Applied Biosystems ABI3130xl Genetic Analyser (Applied Biosystems, UK) was used to resolve the fragments using GeneScan 500-LIZ(−250) as a size standard. Fragment length was established using GeneMapper v5.0 (Applied Biosystems, UK). Genotyping consistency was validated by repeating PCR, fragment analysis and scoring for 10% of samples.

### 2.3 Estimates of genetic diversity

We used Microchecker v2.2.3 (Van Oosterhout et al., 2004) to identify null alleles, allele dropout and stutter peaks, and Bayescan v2.1 (Foll & Gaggiotti, 2008) to test for loci neutrality. GENEPOP v4.2 (Rousset, 2008) was used to test for linkage disequilibrium, deviations from Hardy-Weinberg equilibrium, and to calculate allelic frequencies across populations. GeneAlEx v6.502 (Peakall & Smouse, 2012) was used to assess the number of alleles (*N*_A_), effective alleles (*N_E_*), private alleles (*N*_PA_), expected heterozygosity (*H*_E_), observed heterozygosity (*H*_O_), and to carry out a Mantel test of genetic isolation by distance.

### 2.4 Population genetic structure and patterns of migration

We conducted an Analysis of Molecular Variance (AMOVA) to partition genetic variation at three hierarchical levels (among populations, within populations, and among individuals), and calculated pairwise *F_ST_* values between populations using Arlequin v3.5.2.2. (Excoffier & Lischer, 2010). A Bayesian cluster analysis was conducted in STRUCTURE v2.3.4 (Falush et al., 2007; Hubisz et al. 2009; Pritchard et al., 2000) to estimate the most likely number of genetic clusters (K) informed by individual genotypes. Admixture models with K values ranging from 2 to 15 were considered using twenty iterations, a burn-in length of 10,000 and 50,000 Markov Chain Monte Carlo repeat simulations to quantify the likelihood of each K value. Results were fed into STRUCTURESELECTOR (Li & Liu, 2018) to identify the most likely number of clusters present based on the median of means (MedMeaK), maximum of means (MaxMeaK), median of medians (MedMedK) and maximum of medians (MaxMedK) criteria (Puechmaille, 2016). Bayesian cluster analysis informed by spatial data was conducted using TESS v2.3.1 (Chen et al., 2007) to better understand the extent of spatial genetic structure. Admixture models were run with 50,000 total sweeps, 10,000 burn-in sweeps, and 200 runs per *K*_max_ ranging from 2 to 15. The average Deviance Information Criterion (DIC) of the lowest 10 DIC values was calculated for each *K*_max_ to assess the most likely number of clusters. Runs within 10% of the lowest (DIC) for a given *K*_max_ were used for analysis. CLUMMP v1.1.2 (Jakobsson & Rosenberg, 2007) was used to average variation between repeated iterations for the most likely K values, and the resulting output was visualised using DISTRUCT v1.1.1 (Rosenberg, 2004). A neighbour joining tree assessing the genetic distance of populations was constructed with Populations v1.2.32 (Langella, 2002) using Nei’s standard genetic distance with 1,000 bootstraps per locus and the resulting tree was visualised using TreeView (Page, 2003). The effective number of migrants (*N*_m_) and extent of asymmetrical migration were calculated using div-Migrate (Sundqvist et al., 2016). A matrix was created using 5,000 bootstrap iterations under three alpha values to simulate high (α = 0.05), moderate (α = 0.005) and low (α = 0.0025) levels of gene flow.

### 2.5 Effective population size, population expansion and evidence of genetic bottlenecks

Estimates of effective population size (*N*_e_) for sites containing at least 19 individuals were calculated using the Linkage Disequilibrium Model (LDM) with a critical value of 0.02 in NeEstimator v2.1 (Do et al., 2014). Evidence of population expansion was assessed through the *k* intralocus and *g* interlocus tests (Reich et al., 1998), using the application provided in Bilgin (2007). Evidence of genetic bottlenecks was evaluated in Bottleneck v2.1 (Cornuet & Luikart, 1996) using 1,000 replicates and according to the Two-Phase (TPM) and the Stepwise (SMM) Mutation Models.

### 2.6 Phenotypic variation

Variation in the length-weight relationship between regions (West Atlantic, n = 30; East Atlantic, n = 65; English Channel, n = 60; Baltic Sea, n = 40), was examined by regression analysis on log-transformed data. We calculated relative weight (Wr) as the ratio of the observed weight divided by the predicted weight (from the regression obtained above) to obtain an index of body condition that is more appropriate for fish like lumpfish that have an unusual body shape (Nahdi et al., 2016). The most plausible number of age classes represented in the samples, and the mean size at age (Macdonald & Pitcher, 1979) were calculated through mixture analysis of length-frequency data using PAST v3.17 (Hammer et al, 2001). The Von Bertalanffy growth equation (Kirkwood, 1983) was fitted to estimate growth parameters in each region.

## 3 RESULTS

### 3.1 Population genetic diversity

All microsatellites exhibited polymorphism. The mean number of alleles (*N_A_*) ranged from 4.5 (Ro) to 6.8 (Kl, VB), mean expected heterozygosity (*H_E_*) ranged from 0.592 (Öl) to 0.700 (Kl), and mean *F_IS_* varied from −0.056 (Ro) to 0.110 (Öl) across all loci (Table 1). Initial analysis suggested that null alleles might be present at multiple loci (*Clu34, Clu36, Clu12, Clu33, Clu37* and *Clu40*, Table S2). However, repeatedly removing each locus in turn showed little variation in *F_ST_* values (Tables S3-S8), and therefore all markers were retained for further analyses. No evidence of departures from neutrality or linkage disequilibrium was found after Bonferroni corrections for multiple tests (Rice, 1989). Deviations from Hardy-Weinberg equilibrium (HWE) were detected at 5 of the 15 sites (Table 1), but these involved only 12% of loci after Bonferroni correction (Table S9). The mean number of private alleles (*N_PA_*) was relatively low, ranging from 0.00 to 0.40, with sites in the West Atlantic (FB = 0.30, WB = 0.40) and Baltic Sea (GS = 0.30) showing the highest values.

### 3.2 Population structure and gene flow

Global *F_ST_* was 0.095 (*P* < 0.001) indicating a moderate but significant degree of genetic differentiation. Results of AMOVA indicated that 83.5% of molecular variation was due to variation within individuals, 7% amongst individuals within populations, and 9.5% amongst populations. Pairwise *F_ST_* showed a significant level of genetic differentiation across most populations (Table 2), but populations closer together were genetically similar after Bonferroni correction. On the basis of *F_ST_* values, the strongest differentiation was found between West Atlantic and Baltic Sea populations. Results of a Mantel test support the existence of a significant, albeit weak, isolation by distance (*R*^2^ = 0.1229, *P* = 0.01). The most likely number of genetically distinct groups (*K*) ranged from *K* = 5 (MedMedK, MedMeaK) to *K* = 6 (MaxMedK, MaxMeaK) using STRUCTURESELECTOR (Figure S1). Spatial cluster analysis using TESS suggested a *K_max_* = 10 (Figure S2), though only six of these genetic groups showed substantial representation, and four groups contributed only 3.3% to the genetic background. Distinct clusters were detected in the West Atlantic and Baltic Sea by both STRUCTURE and TESS, with a greater level of admixture across the East Atlantic (Figure 1). Results were consistent in attributing a genetically unique pattern to the Mid Atlantic, English Channel clusters and a Norwegian site at Averøy. A neighbourhood joining tree (Figure 2) showed similar patterns to that of the structuring analyses, highlighting the separation between the West Atlantic and Baltic Sea populations, and the higher degree of admixture within the East Atlantic group.

**Table 2.** Pairwise *F_ST_* values (lower) and Bonferroni adjusted *P* values (upper; Bonferroni correction *P <* 0.00022) between 15 study populations of lumpfish distributed across the natural range of the species using 10 microsatellite loci.

|  | FB | CB | WB | Ha | KI | VB | OH | We | Gu | Na | Av | Ro | KB | Öl | GS |
| --- | --- | --- | --- | --- | --- | --- | --- | --- | --- | --- | --- | --- | --- | --- | --- |
| FB |  | 0.036 | 0.000 | 0.000 | 0.000 | 0.000 | 0.000 | 0.000 | 0.000 | 0.000 | 0.000 | 0.000 | 0.000 | 0.000 | 0.000 |
| CB | 0.013 |  | 0.000 | 0.000 | 0.000 | 0.000 | 0.000 | 0.000 | 0.000 | 0.000 | 0.000 | 0.000 | 0.000 | 0.000 | 0.000 |
| WB | 0.030 | 0.030 |  | 0.000 | 0.000 | 0.000 | 0.000 | 0.000 | 0.000 | 0.000 | 0.000 | 0.000 | 0.000 | 0.000 | 0.000 |
| Ha | 0.130 | 0.112 | 0.117 |  | 0.000 | 0.000 | 0.000 | 0.000 | 0.000 | 0.000 | 0.000 | 0.000 | 0.000 | 0.000 | 0.000 |
| KI | 0.120 | 0.101 | 0.098 | 0.050 |  | 0.081 | 0.000 | 0.000 | 0.000 | 0.000 | 0.000 | 0.000 | 0.000 | 0.000 | 0.000 |
| VB | 0.117 | 0.093 | 0.102 | 0.042 | 0.011 |  | 0.000 | 0.000 | 0.000 | 0.018 | 0.000 | 0.000 | 0.243 | 0.000 | 0.000 |
| OH | 0.152 | 0.111 | 0.124 | 0.049 | 0.034 | 0.021 |  | 0.000 | 0.009 | 0.252 | 0.000 | 0.000 | 0.000 | 0.000 | 0.000 |
| We | 0.177 | 0.154 | 0.146 | 0.065 | 0.056 | 0.042 | 0.024 |  | 0.324 | 0.000 | 0.000 | 0.000 | 0.000 | 0.000 | 0.000 |
| Gu | 0.188 | 0.160 | 0.152 | 0.083 | 0.060 | 0.057 | 0.014 | 0.003 |  | 0.009 | 0.000 | 0.000 | 0.000 | 0.000 | 0.000 |
| Na | 0.157 | 0.121 | 0.128 | 0.080 | 0.035 | 0.020 | 0.004 | 0.029 | 0.029 |  | 0.036 | 0.000 | 0.018 | 0.000 | 0.000 |
| Av | 0.153 | 0.122 | 0.108 | 0.102 | 0.027 | 0.021 | 0.042 | 0.061 | 0.061 | 0.018 |  | 0.000 | 0.000 | 0.000 | 0.000 |
| Ro | 0.142 | 0.132 | 0.138 | 0.057 | 0.039 | 0.041 | 0.043 | 0.065 | 0.061 | 0.048 | 0.075 |  | 0.000 | 0.000 | 0.000 |
| KB | 0.113 | 0.085 | 0.095 | 0.034 | 0.021 | 0.004 | 0.028 | 0.048 | 0.067 | 0.022 | 0.037 | 0.046 |  | 0.000 | 0.000 |
| Öl | 0.194 | 0.139 | 0.176 | 0.087 | 0.129 | 0.115 | 0.097 | 0.105 | 0.113 | 0.126 | 0.152 | 0.149 | 0.088 |  | 0.216 |
| GS | 0.187 | 0.154 | 0.181 | 0.097 | 0.140 | 0.132 | 0.134 | 0.136 | 0.152 | 0.159 | 0.175 | 0.152 | 0.110 | 0.011 |  |

**Figure 1.**
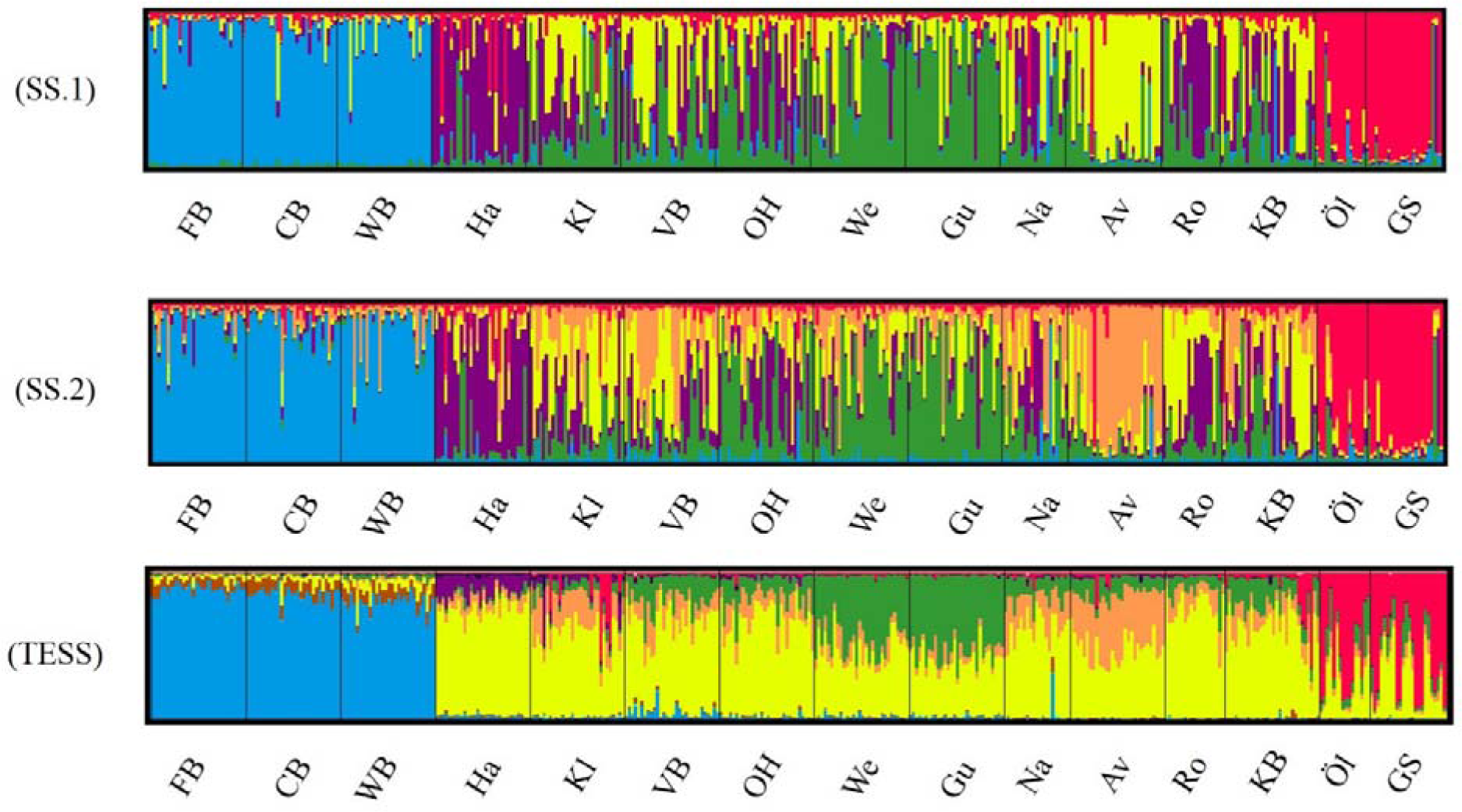
Lumpfish genetic structuring according to STRUCTURESELECTOR with (a) MedMedK and MedMeanK of *K* = 5, (b) MaxMedK and MaxMeanK of *K*= 6, and (c) TESS with *K*_max_ = 10 based on lowest mean DIC value. Each bar represents one individual with colours indicating probability of belonging to different genetically distinct groups.

**Figure 2.**
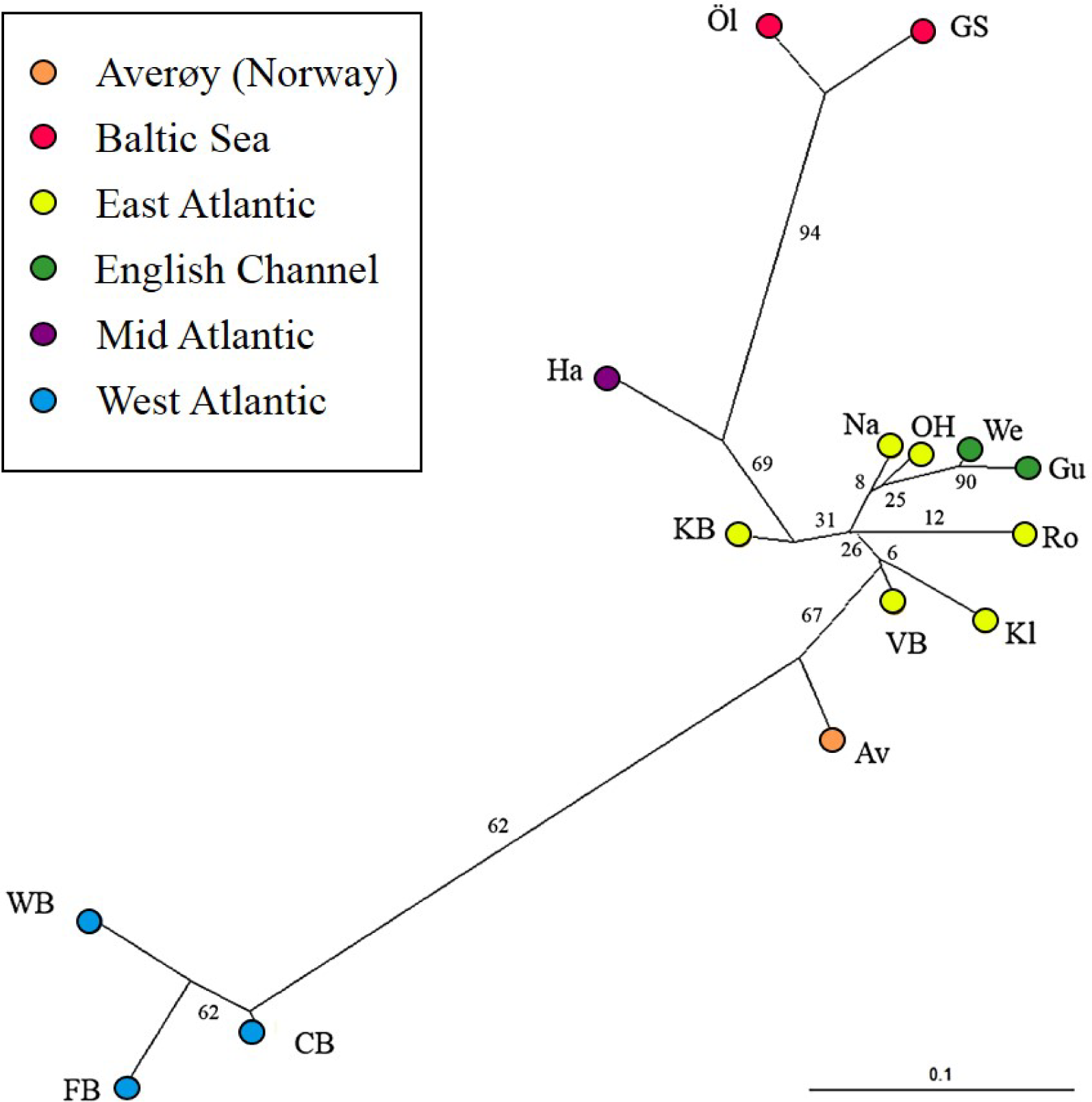
Neighbourhood joining tree (based on Nei’s Standard Genetic Distance) of 15 lumpfish populations genotyped with 10 microsatellite loci.

The effective number of migrants (*N*_m_) ranged from 1.00 between sites in the English Channel to 0.03 between sites in the West Atlantic and Baltic Sea. The exchange of migrants was much higher within genetic clusters than among clusters (Table S10), with the highest levels of gene flow found within the East Atlantic and within the English Channel (Figure 3). The only evidence of moderate asymmetric gene flow was from Norway towards the Faroe Islands (*N*_m_ = 0.507), but this was only detected at α = 0.05, and not at the lower α-values (0.005 or 0.0025).

**Figure 3.**
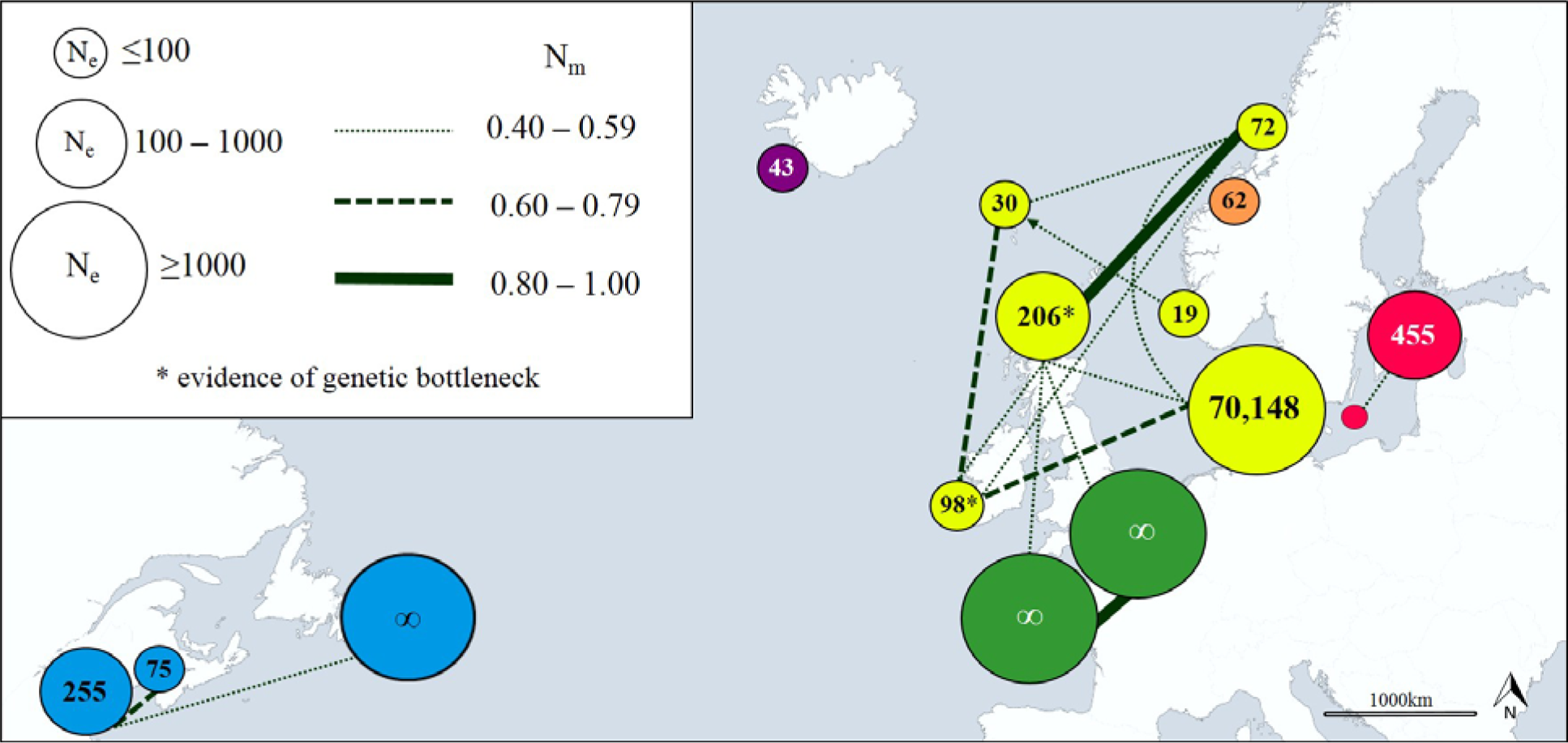
Patterns of gene flow among lumpfish populations with colours indicating genetic groups, symbol size proportional to effective population size (*N*_e_), line thickness proportional to effective number of migrants (*N*_m_), and line direction indicative of significant asymmetric gene flow.

### 3.3 Effective population size, population expansion and evidence of genetic bottlenecks

Estimates of effective population size (*N*_e_) based on a Linkage Disequilibrium Model (LDM) varied from 19 (Norway) to 70,148 (Denmark; Table S11). Sites with low *N*_e_ values (<75) were found across Iceland, Faroe Islands and Norway (Figure 3). No evidence of recent population expansions was found according to the intra-locus *k* and inter-locus *g* tests (Table S12). A significant deficiency of heterozygotes was identified in Ireland and Scotland using the Single Mutation Model (SMM) in BOTTLENECK (Wilcoxon signed-rank test, *P* = 0.0033 after Bonferroni correction), suggesting that these populations could have undergone a recent genetic bottleneck (Table S13), but this was not detected by the Two-Phase Model of Mutation (TPM).

### 3.4 Phenotypic variation

The relationship between length and weight differed significantly between regions (*F*_4,192_ = 917.2, *P* < 0.001; Figure 4). Lumpfish in the Baltic Sea were heavier relative to their size than lumpfish in the East Atlantic and the English Channel (pairwise comparisons: Baltic - East Atlantic, estimate = −0.090 ± 0.036, t = −2.530, *P* = 0.012; Baltic - English Channel, estimate = −0.145 ± 0.046, *t* = −3.171, *P* = 0.002), but were similar to those in the West Atlantic (pairwise comparison Baltic - West Atlantic, estimate = −0.094 ± 0.050, *t* = −1.891, *P*= 0.060). The relative weight of lumpfish differed between regions (*F*_3,191_ = 2.841, *P* = 0.039) and was highest in the Baltic Sea and the West Atlantic, and lowest in the East Atlantic and the English Channel.

**Figure 4.**
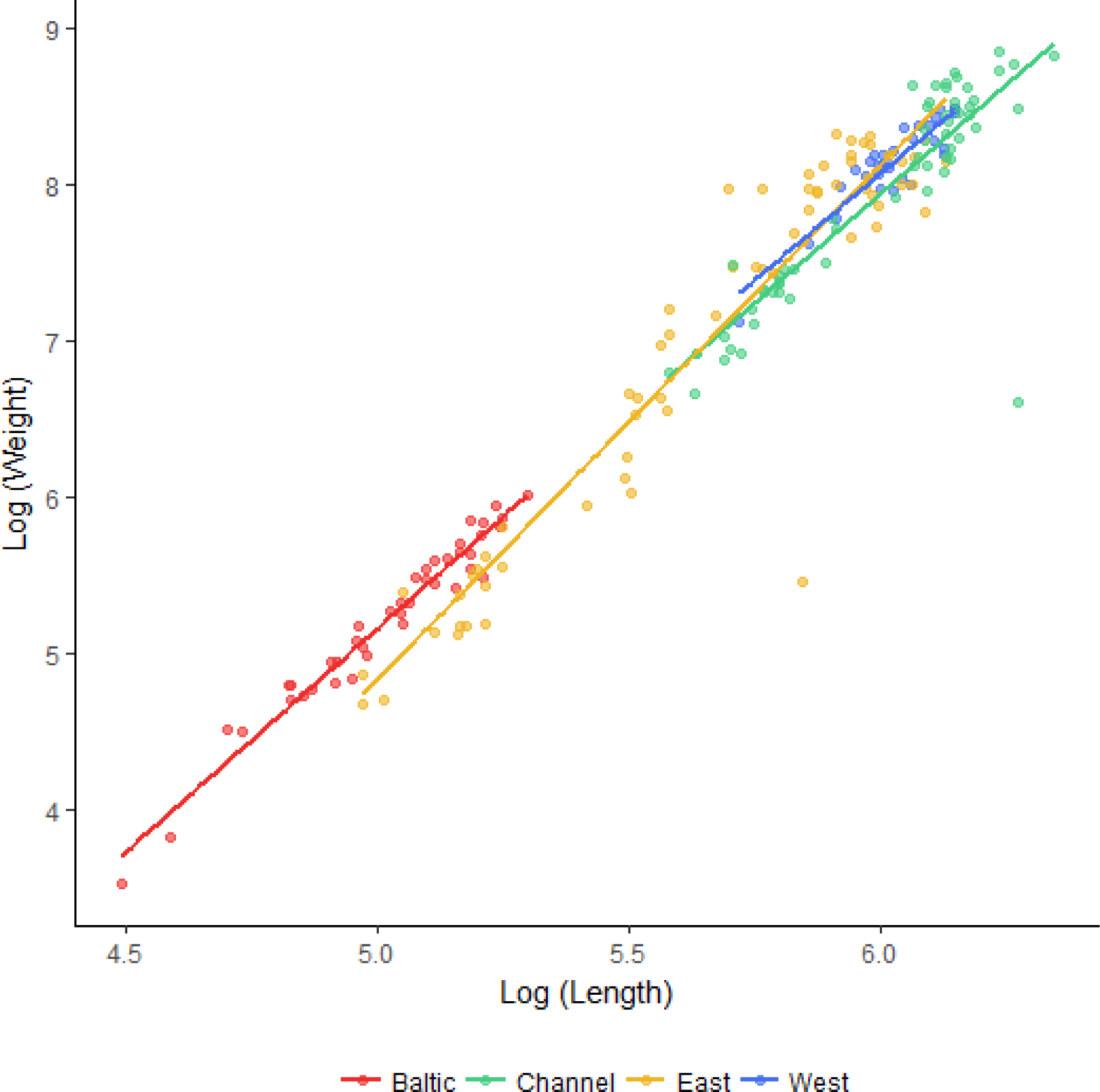
Length-weight relationships (log_10_ scale) for lumpfish sampled in the Baltic Sea, English Channel, East Atlantic and West Atlantic.

Mixture analysis identified multiple plausible age classes present amongst lumpfish sampled in the Baltic Sea (7 age classes), East Atlantic (4 age classes) and English Channel (3 age classes), but only a single plausible age class in the West Atlantic. Based on the parameters of the Von Bertalanffy Growth equation, the maximum age was estimated to be 6.0 years for Baltic populations, 5.7 yrs for populations in the East Atlantic and 7.5 yrs for southern populations spawning in the English Channel. Fitted growth equations differed significantly between regions (Table 3), with lumpfish in the Baltic Sea showing the slowest growth and those in the English Channel showing the fastest.

**Table 3.** Von Bertalanffy growth parameters (*L*∞ = asymptotic length, *t*_0_ = initial condition parameter, and *K* = Brody growth rate or curvature parameter) and estimated mean weight at first maturity (± 95 CI) for lumpfish from different genetically distinct regions.

| Region | Von Bertalanffy Growth parameters |  |  | Weight<br>at 1<sup>st</sup> maturity (g) |
| --- | --- | --- | --- | --- |
| | $L_{\infty}$ (mm) | $t_0$ | $K$ (yr<sup>-1</sup>) | |
| Baltic Sea | 200 ± 6 | 0.14 ± 0.02 | 0.51 ± 0.02 | 150 ± 12.5 |
| East Atlantic | 461 ± 14 | 0.36 ± 0.23 | 0.56 ± 0.09 | 2,019 ± 265.5 |
| English Channel | 571 ± 22 | -1.08 ± 0.59 | 0.35 ± 0.20 | 3,007 ± 519.5 |

## 4 DISCUSSION

Our study reveals a significant degree of structuring in lumpfish populations which is consistent with moderate isolation by distance, and should inform the translocation of lumpfish across salmon farms. Genetically distinct groups were found in the West Atlantic (USA, Canada), Mid Atlantic (Iceland), East Atlantic (Faroe Islands, Ireland, Scotland, Norway, Denmark), English Channel (England), Averøy (Norway) and Baltic Sea (Sweden). Whilst significant gene flow was detected within each of these groups, little exchange of migrants was found between these areas. Our results also indicate the existence of significant phenotypic differences across the range, that mimic to some extent the observed genetic differences. Thus, lumpfish from the Baltic Sea were not only genetically distinct, they were also found to be smaller, grow at a slower rate and weigh more relative to their size than lumpfish from the North Atlantic. Although our growth estimates were based on length frequency data and did do not distinguish between males and females, they are in line with estimates based on mark and recapture studies in Norway and Iceland (*L*∞ = 527 ± 64 mm, *K*= 0⋅26 ± 0⋅14 year^−1^; Kasper et al., 2014), and suggest that Baltic lumpfish grow more slowly and mature at a much smaller size (c. 150 g) than lumpfish from the North Atlantic (2.0-3.0 Kg). The slow growth shown by Baltic lumpfish may be of interest for selective breeding programmes in aquaculture, as slow growing cleaner fish may be better suited for feeding on sea lice (Powell et al., 2017).

Pampoulie et al. (2014) first suggested that lumpfish in the West and East Atlantic were separated by cold southward polar currents, and that populations in the Baltic Sea may have become isolated during the Last Glacial Maximum. Though our analysis supports this broad division, it also indicates a finer population structure, revealing that lumpfish in the mid Atlantic and English Channel are genetically distinct from other populations. The conclusion of our genetic analyses is consistent with recent tagging studies in Norway and Iceland showing that whilst lumpfish can move offshore to feed, they return to spawn in their home waters (Kennedy et al., 2015; Kennedy et al., 2016) and do not migrate between Iceland and Norway (Kasper et al., 2014). There is little information on southern lumpfish populations, though lumpfish in the English Channel appear to spawn earlier in the season than populations further north (Powell et al., 2018), probably due the warmer temperatures and better feeding opportunities, which are known to influence maturation and spawning of lumpfish (Hedeholm et al., 2017). It is thus possible that the warmer waters found at the species’ southern range may favour an early spawning and lead to some degree of reproductive isolation, hence limiting gene flow along a latitudinal gradient. With the exception of the Averøy population, the remaining sites in the East Atlantic appear to be genetically uniform, as reported along the Norwegian coast (Jónsdóttir et al., 2017).

The level of genetic diversity, and therefore the ability to adapt and respond to selection, differed substantially among regions. Our estimates of effective population size, the first for this species, were particularly low across the North East Atlantic (Iceland, *N*_e_ = 43; Faroe Islands, *N*_e_ = 30; Norway, mean *N*_e_ = 51), and some evidence of genetic bottlenecks was also detected at sites in Ireland and Scotland, though the evidence for this was not strong. The North East Atlantic supports the largest lumpfish roe fishery (Jónsdóttir et al., 2018), with a production of 4,000 tonnes of roe per year (Johannesson, 2006). Given a maximum yield of c. 4kg roe/female (Johannesson, 2006), this level of harvest likely surpasses 1 million mature females every year. Harvesting for lumpfish roe is both size and sex-selective, which increases the vulnerability of populations to over-exploitation (Hoenig & Hewitt, 2005; Ratner & Lande, 2001) and may explain the low estimates of effective population size found across this area. The North East Atlantic populations appear to be small and reducing pressure on these stocks would decrease the risk of over exploitation.

### 4.1 Conclusions and management implications

By 2020 c. 50 million lumpfish will be required by the salmon farming industry (Powell et al., 2017; Treasurer, 2018) and most of these will come from the stripping of wild broodstock (Wittwer & Treasurer, 2018) caught in Iceland and Norway, and then shipped as eggs or larvae to salmon farms elsewhere. Information on lumpfish escapees is lacking but corkwing wrasse (*Symphodus melops*) deployed as cleaner fish in Norway have recently been found to escape and hybridise with local populations (Faust et al., 2018), and the same could happen with lumpfish. Efforts should thus be made to reduce the risk of lumpfish escaping from fish farms and interbreeding with local populations, as high propagule pressure associated with open-net pens is the single most important factor determining the impact of escapees (Consuegra et al., 2011).

Our study suggests that lumpfish translocations should be restricted within genetically homogenous groups to reduce the risk of genetic introgression between native and non-native populations. In this sense, lumpfish from some areas of Norway, and particularly from Iceland, may be ill-suited for deployment in Ireland, Scotland and the Faroe Islands, and vice-versa. Ultimately, closing the breeding cycle of the species in captivity, and producing sterile lumpfish for deployment in salmon farms, must be a research priority for both the conservation of the species and the cleaner fish industry (Powell et al., 2017), as this will lessen dependency on wild broodstock and reduce the risk of genetic introgression.

## AKNOWLEDGMENTS

This work was partially funded by Marine Harvest Scotland through the LUMPFISH project and the Welsh Government via the European Regional Development Fund (SMARTAQUA Operation). We are indebted to Majbritt Bolton-Warberg, Danny Boyce, Werner Forster, Gus Galloway, Ása Johannesen, Lars Jørgen Ulvan, Niklas Larson, Marine Harvest Scotland, Michael Pietrak, Adam Rainsden and Peter Rask for supplying tissue samples. We are also grateful to Chloe Robinson, Niall Coates, Christine Gray, Craig Pooley and Ian Tew for assistance in optimising primers and processing samples.

## AUTHORS’ CONTRIBUTIONS

CGL and SC designed the study, wrote the project and secured the funding. BAW processed the samples and carried out the genetic analysis with assistance from SC. BAW and CGL wrote the MS with input from SC. BAW and CGL carried out the statistical analysis. All authors approved the final submission.

